# Wasting is essential: cachexia is driven by a wound-healing adaptation

**DOI:** 10.64898/2026.10.23.761459

**Authors:** Cameron Haynes, Christopher Whelan, Joel Brown

## Abstract

**Background:** Several explanations have been given for why cachexia occurs in the presence of severe acute wounds and chronic diseases. None of these explanations address the scope of clinical settings in which cachexia has been observed, nor do they account for the failure of nutritional support to reverse cachexia. We hypothesize that induced muscle wasting (IMW) is an adaptation to support wound healing, and when a wound does not heal, prolonged IMW leads to cachexia.

**Methods:** To evaluate this hypothesis, we developed a mathematical model of IMW. In this model, we compared the time needed for wound healing when IMW was present and when it was inhibited. We also measured the relationship between the pathogen growth rate and final patient muscle mass for chronic diseases such as cancer and tuberculosis, demonstrating the onset of cachexia. Finally, we simulated the effects of nutritional support and contrasted the outcomes of when IMW was present and when it was inhibited.

**Results:** Acute wound healing was ∼2-6× longer without IMW than with IMW. For chronic wounds, steady-state muscle mass strongly depended on the growth rate of the pathogenic agent. Inhibiting IMW was more effective at preventing acute muscle mass loss than nutritional support, and the benefit of nutritional support was enhanced by inhibiting IMW. Nutritional support also increased the speed of healing acute wounds when IMW was inhibited. For cancer with high growth rate, giving nutritional support alone had a negligible effect on steady-state muscle mass. But when IMW was inhibited, muscle mass loss was attenuated. With IMW inhibition, nutritional support of 1× normal diet was able to abrogate muscle mass loss entirely.

**Conclusions:** Inhibiting IMW slows the healing of acute wounds, supporting our hypothesis that IMW is an adaptation to support rapid wound healing. Our experiments demonstrate that nutritional support is insufficient to prevent severe muscle mass loss while IMW continues. But when IMW is inhibited, nutritional support restores normal levels of muscle mass for both acute and chronic wounds, as well as increasing the rate of wound healing. Discovering and developing methods to inhibit IMW for clinical use will be essential to restore patient muscle mass, ease suffering, and reduce mortality.

## 1. Introduction

Cachexia is a wasting condition known to accompany many different states of injury and disease (hereafter referred to as *conditions*), including burns, chronic obstructive pulmonary disorder (COPD), HIV, tuberculosis (TB), and cancer [1]. The hallmark of cachexia is muscle loss that often coincides with fat wasting, but not always [2]. This progressive atrophy increases functional impairment and suffering for patients, eventually leading to death if the underlying pathology is not resolved. Despite the ubiquity of cachexia across diverse chronic diseases, no FDA-approved treatments are available for the condition.

Efforts to reverse cachexia have been stymied by its complex etiology. Decades of fruitful research into its molecular mechanisms show that muscle and fat wasting are induced by a host of bloodborne factors. Experiments have demonstrated that cachexia can be transmitted from one animal to another via plasma transfusions [3]. These factors (i.e., cytokines, glucocorticoids, etc.) are abundantly generated during the course of any disease [4]. Thus, cachexia results from the prolonged exposure of muscle and fat tissue to signals capable of inducing wasting. However, a crucial question remains unanswered. Why are these tissues, and muscles in particular, primed for this induced response in the first place?

In cancer research, several explanations have been posited. First, some researchers argue that muscle wasting results from an excessive energy demand placed on the body by the tumor, and thus muscle wasting is akin to starvation [5]. However, this does not accord with the fact that fat does not always waste during cachexia, nor does it explain the experiments in which bloodborne factors can cause cachexia in otherwise healthy controls. Second, other researchers have proposed that a tumor secretes wasting-inducing factors to nourish itself. Though initially plausible, this hypothesis does not account for the general appearance of cachexia in unresolved chronic diseases, especially in those conditions not caused by a pathogen, such as COPD or chronic heart failure [6]. This paradigmatic problem is accentuated by the routine observation of cachexia in patients with severe burns. A different explanation is in order – one that encompasses cachexia from all maladies.

Some of the bloodborne factors responsible for inducing muscle wasting during cachexia also influence other bodily components. For example, IL-6—a well-known cytokine produced in all the conditions mentioned above —induces muscle atrophy, but it also induces the hepatic acute-phase response (APR) [7–8]. During APR, hepatocytes use large quantities of amino acids to produce proteins essential for a vigorous immune response (known as *acute-phase proteins*) [8]. Therefore, IL-6 simultaneously causes release of amino acids from the muscles and their use in protein synthesis in the liver. Might these responses be mechanistically connected?

We propose that induced muscle-wasting (IMW) in response to pathogens and injury is an *adaptation* of the human body: inflammatory signals cause muscles to release amino acids to supply the demands of the hepatic APR and other subsequent wound-healing efforts. Thus, IMW can benefit wound healing in acute contexts, in which the duration of the wasting is limited. However, if a wound does not heal rapidly, this response is prolonged, culminating in profound atrophy.

The historical support for our paradigm begins with Sir David Cuthbertson and his initial inquiries into the metabolic changes resulting from injury. He demonstrated that the stoichiometric ratios of substances excreted in patients’ urine indicated the muscle catabolism. He further proposed that breaking down muscle supported the “exigencies of [wound] repair [9–10].” Reeds et al., in an article examining the fate of amino acids released from muscle during injury, concluded that nitrogen-loss data were well-explained if most of the amino acids were used to support the hepatic APR, with a minority catabolized for gluconeogenesis [11]. And the late Kenneth Fearon noted that muscle wasting in cancer could be explained through a connection to the injury response, in which cancer cachexia emerged as a “double-edged sword” of naturally-occurring host mechanisms related to wound repair [6]. Our *adaptation paradigm* synthesizes these perspectives. The paradigm clarifies the complex phenomena of cachexia, and it offers a testable hypothesis: if IMW supports wound healing, then when it is inhibited, wound healing should be impaired.

Here, we develop a mathematical model of cachexia to evaluate this hypothesis. After demonstrating that IMW supports wound healing in an acute-wound context (i.e., healing from a burn), we consider the model’s predictions for two chronic disease settings. Finally, we use our model to demonstrate that without inhibiting IMW, nutritional support fails to reverse cachexia, but when IMW is inhibited, nutritional support recapitulates its supportive effect on wound healing and maintains muscle mass.

## 2. Methods

### 2.1 The Necessity of Mathematical Modeling

The goal of this study is to evaluate the effect of IMW on wound healing. To approach such an objective in a wet-laboratory setting would require systematically inhibiting IMW without affecting other physiological processes. To the best of our knowledge, this is currently not feasible; otherwise, cachexia could be clinically managed through such a technique.

Mathematical modeling solves this impasse. Through recapitulating the essential physiological phenomena of cachexia, such a model allows for a qualitative assessment of our hypothesis, because it is comparatively easy to adjust IMW as an independent variable, leaving other values fixed. Therefore, we construct a mathematical model that retains the necessary components of IMW, while avoiding unnecessary detail.

### 2.2 Prior Mathematical Models of Cachexia

Few mathematical models of cachexia exist in the literature, and most embrace one of the two explanations for cachexia refuted above (Table 1). Assuming that muscle wasting results from the severe caloric imbalance caused by a tumor, some have modeled cachexia using the dynamic energy budget theory that chiefly considers the nutritional needs of the patient [5, 12]. Others have modeled the effect of tumor volume on muscle size and the satellite-cell pool [13]. Cancer is by far the most common cause of cachexia used in modeling studies. However, some focus on the effects of cancer treatment on wasting, concluding that the second-order effects of treatment should be considered for appropriate patient care [14]. In contrast, instead of seeking a direct application to therapeutic efforts, Kobak et al. used mathematical modeling of muscle wasting to challenge the efficacy of certain techniques commonly used to measure fractional protein synthesis in the cachexia field [15]. Our model differs from these previous studies, in mathematical form, scope, and goals.

**Table 1.**
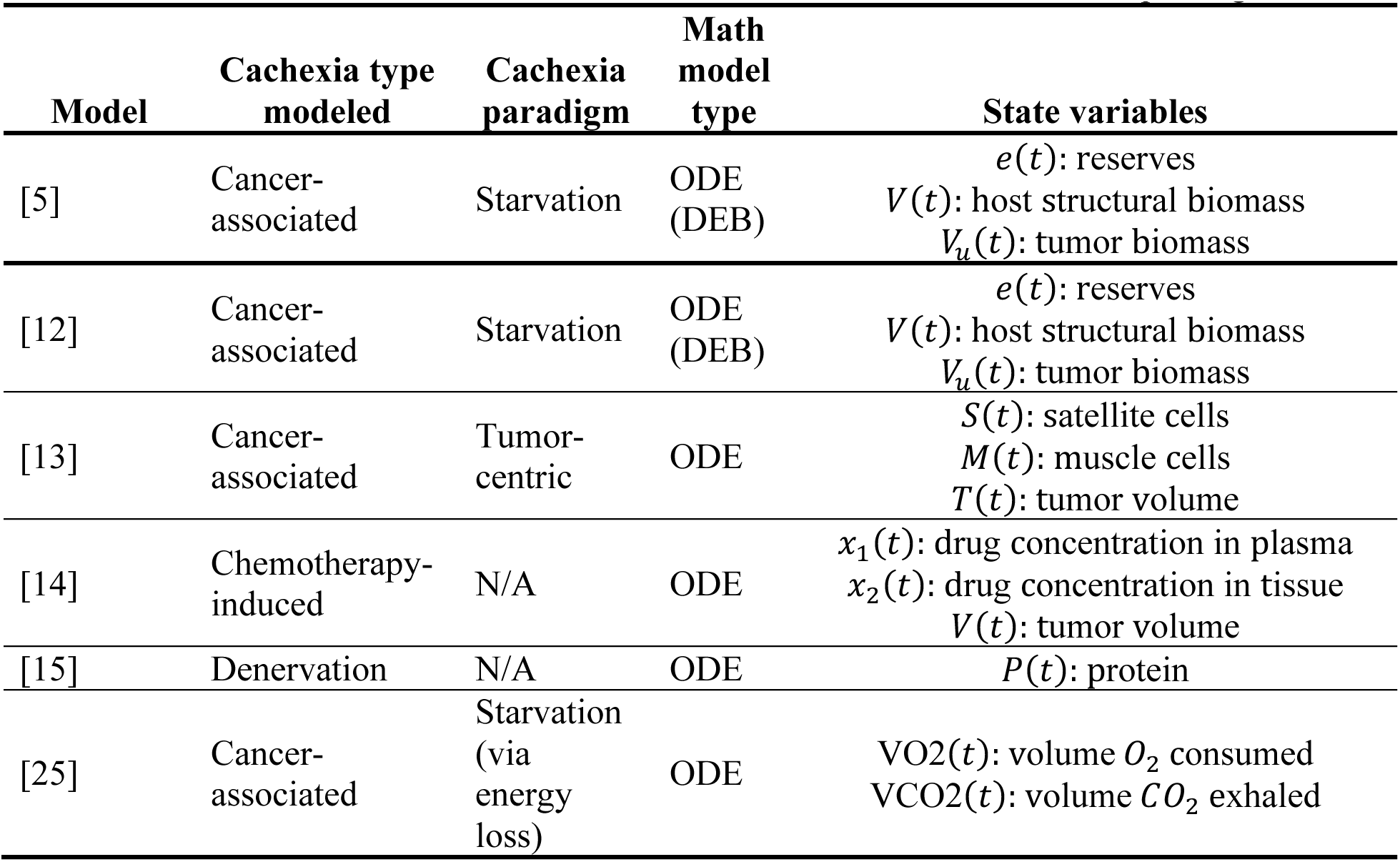
Mathematical models of cachexia feature starvation or tumor-centric paradigms.

### 2.3 Our Model

We modeled cachexia using a biochemical reaction network that includes the major physiological components of IMW: the wound, muscle, and the liver (Figure 1). The behavior of each of these components and their interactions with bloodborne mediators are represented by biochemical reaction equations, and we applied mass action kinetics to these reactions to derive an ordinary differential equation for simulating the system in time (Table 2).

**Figure 1.**
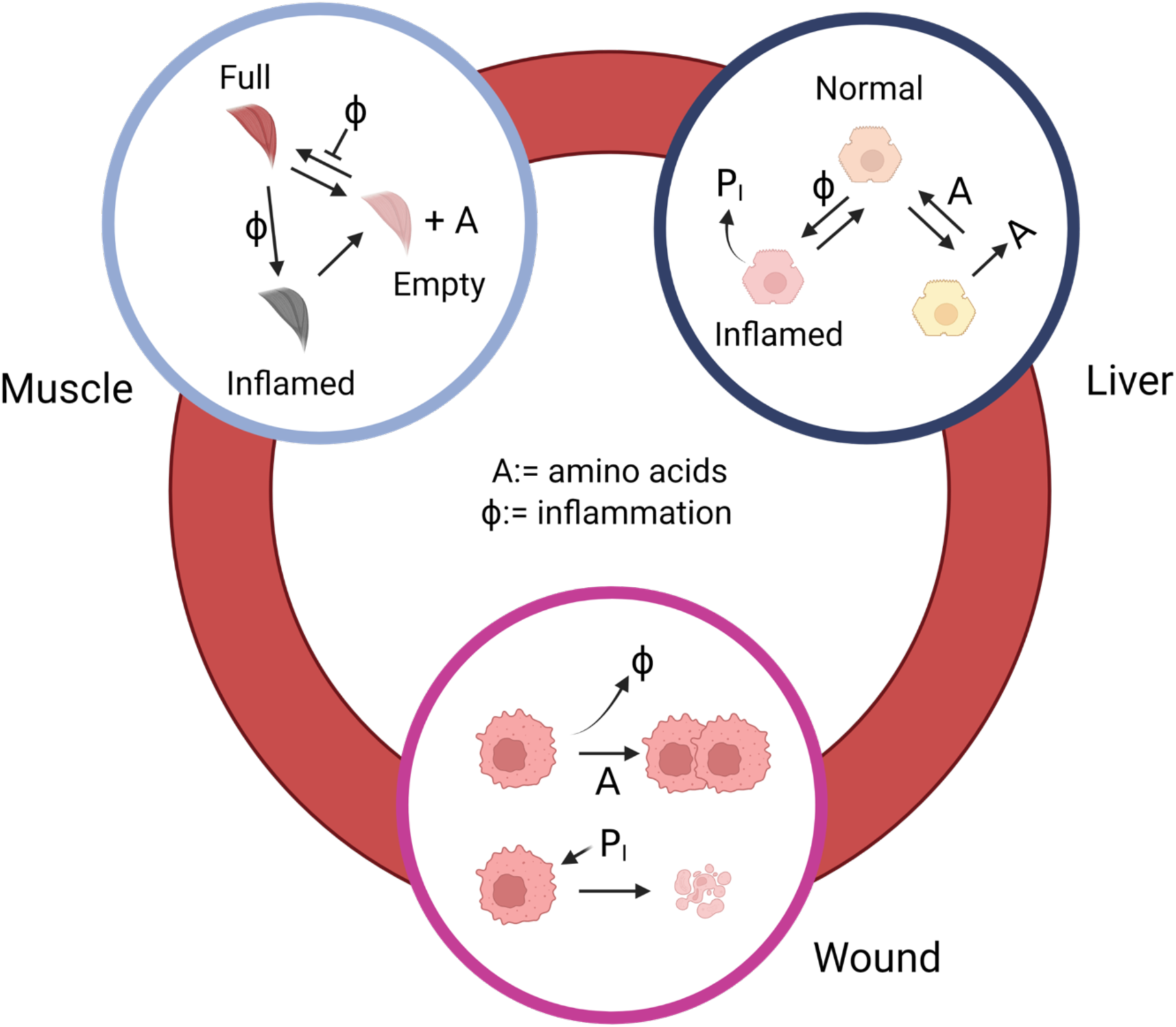
The compartments of the ODE Model and their interactions. Note that the muscle, naturally in a *full* state, can undergo proteolysis in response to either low amino acid levels (A) or inflammation (*φ*) and enter the *empty* state. This is the basis of muscle wasting in the model. These nutrient levels then affect the function of the liver and the rate at which it can produce immune proteins (*P_I_*), thereby driving the rate of wound clearance.

**Table 2.** A simple mathematical model, derived from mass action kinetics, recapitulates cachexia physiology.

| Model component | System variables | Differential equations |
| --- | --- | --- |
| Liver | $L_N$ = resting hepatocytes<br>$L_A$ = amino acid generating hepatocytes<br>$L_\phi$ := inflamed (APR) hepatocytes | $\dot{L}_N = -k_1 L_N + k_3 L_A A - k_{13} L_N \phi + k_{15} L_\phi$<br>$\dot{L}_A = k_1 L_N - k_3 L_A A - k_{14} L_A \phi$<br>$\dot{L}_\phi = k_{13} L_N \phi + k_{14} L_A \phi - k_{15} L_\phi$ |
| Muscle | $M_f$: full myocytes<br>$M_e$: empty myocytes<br>$M_\phi$: full inflamed myocytes<br>$M_{e\phi}$: empty inflamed myocytes | $\dot{M}_f = -k_4 M_f + k_5 M_e A - (k_7 + k_{21}) M_f \phi + k_{10} M_\phi$<br>$\dot{M}_e = k_4 M_f - k_5 M_e A - (k_8 + k_{22}) M_e \phi + k_{11} M_{e\phi}$<br>$\dot{M}_\phi = k_7 M_f \phi - (k_9 + k_{10}) M_\phi$<br>$\dot{M}_{e\phi} = k_8 M_e \phi + k_9 M_\phi - k_{11} M_{e\phi}$ |
| Wound | $W$: wound cells<br>$C$: cancer cells<br>$TB$: cells infected with *Mycobacterium tuberculosis* | $\dot{W} = -k_{17} W P_I + 20k_{23} CA + 20k_{26} TB$<br>$\dot{C} = k_{23} CA - k_{25} C P_I$<br>$\dot{TB} = k_{26} TB - k_{28} TBP_I$ |
| Dietary Intake | $D$: active dietary intake<br>$D_\phi$: inactivated dietary intake | $\dot{D} = -k_{19} D \phi + k_{20} D_\phi$<br>$\dot{D}_\phi = k_{19} D \phi - k_{20} D_\phi$ |
| Blood | $A$: amino acid concentration<br>$\phi$: pro-inflammatory molecules<br>$P_I$: acute-phase proteins | $\dot{A} = (k_2 - k_3 A) L_A + k_4 M_f - k_5 M_e A - k_6 M_f A + k_9 M_\phi - k_{16} L_\phi A + k_{18} D + k_{23} CA$<br>$\dot{\phi} = -[(k_7 + k_{21}) M_f + (k_8 + k_{22}) M_e] \phi + k_{12} W + k_{24} C + k_{27} TB - (k_{13} L_N + k_{14} L_A) \phi$<br>$\dot{P}_I = k_{16} L_\phi A - k_{17} W P_I - k_{25} C P_I - k_{28} TBP_I$ |

The liver component comprises three states that hepatocytes can take: (1) *normal* (i.e., resting); (2) *synthesizing* amino acids from bloodborne nutrients; or (3) *APR*, producing acute-phase proteins in response to inflammatory signals. Normal hepatocytes enter into the amino-acid generating state (Reaction 1), where they produce amino acids from other substrates (Reaction 2). Then, they are restored to the normal state by sensing amino acids (Reaction 3).

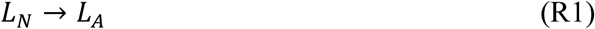

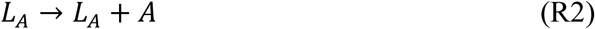

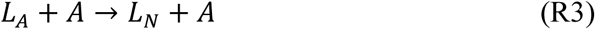

Through this negative feedback loop, the liver component maintains an equilibrium concentration of amino acids during homeostasis.

Without inflammation, the muscle component comprises two states of myocytes—one in which the myocyte has not undergone wasting, called *full* (*M_f_*), and one in which it has broken down many of its contractile proteins and released the amino acids, called *empty* (*M_e_*). Full myocytes enter the empty state and release their amino acids (Reaction 4). Conversely, empty myocytes are restored to being full by interacting with amino acids (Reaction 5).

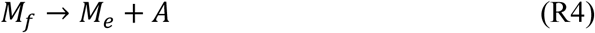

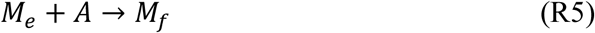

These reactions enable the system, during homeostasis, to maintain an equilibrium muscle mass based on the demand for amino acids. Moreover, the full myocytes, when there is no inflammation, use up a small amount of amino acids in their metabolism (Reaction 6).

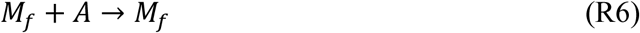

However, when full myocytes receive inflammatory signals, they enter an inflamed state (*M_φ_*) (Reaction 7). Empty myocytes become inflamed similarly (*M_eφ_*) (Reaction 8). Full, inflamed myocytes then release their amino acids into circulation (Reaction 9). They cannot become full again until they become uninflamed again (Reactions 10 or 11, then Reaction 5). This is our model’s implementation of IMW.

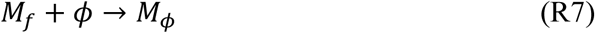

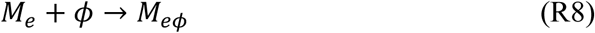

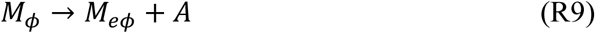

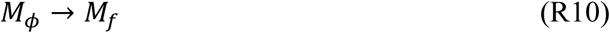

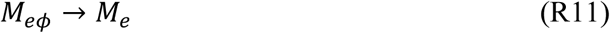

Finally, the wound compartment represents damaged bodily tissue that is actively inflamed, potentially resulting from one of several causes. In the case of a burn, the definition of the wound is straightforward. However, in the case of cancer and TB, the connection between wounding and disease burden is less clear. Therefore, in the case of cancer and TB, we model the wound compartment as proportional to tumor or disease burden, respectively.

The wound interacts with the system in a few crucial ways. By releasing inflammatory cues (Reaction 12), the wound induces the muscle-wasting response (Reaction 7) and APR (Reactions 13-14) in the liver. Then, the APR hepatocytes slowly return to a normal state (Reaction 15).

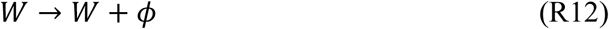

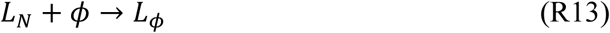

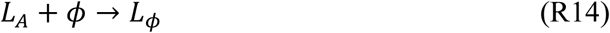

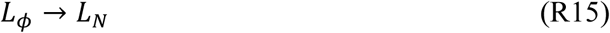

The APR hepatocytes synthesize acute-phase proteins (Reaction 16). These proteins facilitate the immune clearance of wound cells (Reaction 17), decreasing the wound size. Then, the acute-phase proteins are also slowly filtered out of the blood at the liver and other sites (Reaction 18).

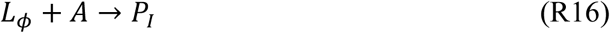

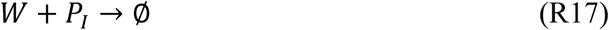

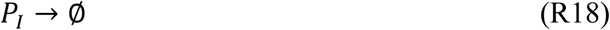

Through these interactions, inducing the muscle wasting response forms a negative feedback loop with the goal of clearing and healing the wound. The wound releases inflammatory signals that activate the APR in the liver and IMW in the muscle. Then, the APR hepatocytes use the increased amino acid availability from IMW to synthesize large numbers of acute-phase proteins (*P_I_*). In turn, these proteins enable the immune system to clear wound cells, reducing the overall output of inflammatory signals.

To parameterize the model, we initialized all of the reaction rate parameters with a value of 1. By adjusting the reaction rates, we qualitatively matched the model output to the typical course of IMW found in mice with severe burns [16]. The final parameter values used can be found in Table 3 and the project’s GitHub repository (see Data and Code Availability).

**Table 3.** Parameter values that qualitatively capture IMW.

| Parameter | Value (units) |
| --- | --- |
| $k_1$ (Rate of Reaction 1) | 1 (per time) |
| $k_2$ (Rate of Reaction 2) | 1 (per time) |
| $k_3$ (Rate of Reaction 3) | 1 (per time) |
| $k_4$ (Rate of Reaction 4) | 1 (per time) |
| $k_5$ (Rate of Reaction 5) | 1 (per time) |
| $k_6$ (Rate of Reaction 6) | 0.015 (per time) |
| $k_7$ (Rate of Reaction 7) | $R$ (See Section 2.7) (per time) |
| $k_8$ (Rate of Reaction 8) | $R$ (See Section 2.7) (per time) |
| $k_9$ (Rate of Reaction 9) | 1 (per time) |
| $k_{10}$ (Rate of Reaction 10) | 0.01 (per time) |
| $k_{11}$ (Rate of Reaction 11) | 0.01 (per time) |
| $k_{12}$ (Rate of Reaction 12) | 1 (per time) |
| $k_{13}$ (Rate of Reaction 13) | 1 (per time) |
| $k_{14}$ (Rate of Reaction 14) | 1 (per time) |
| $k_{15}$ (Rate of Reaction 15) | 1 (per time) |
| $k_{16}$ (Rate of Reaction 16) | 1 (per time) |
| $k_{17}$ (Rate of Reaction 17) | 1 (per time) |
| $k_{18}$ (Rate of Reaction 18) | 1 (per time) |
| $k_{19}$ (Rate of Reaction 19) | 1 (per time) |
| $k_{20}$ (Rate of Reaction 20) | 1 (per time) |
| $k_{21}$ (Rate of Reaction 21) | 1 (per time) |
| $k_{22}$ (Rate of Reaction 22) | $1 - R$ (See Section 2.7) (per time) |
| $k_{23}$ (Rate of Reaction 23) | $1 - R$ (See Section 2.7) (per time) |
| $k_{24}$ (Rate of Reaction 24) | Varied between 0 and 0.01 (per time) |
| $k_{25}$ (Rate of Reaction 25) | Acute Wound: [4, 8, 12]; 8 if cancer or tuberculosis (per time) (See Section |
| $k_{26}$ (Rate of Reaction 26) | 1 (per time) |
| $k_{27}$ (Rate of Reaction 27) | Varied between 0 and 0.01 (per time) |
| $k_{28}$ (Rate of Reaction 28) | 8 (per time) |
| $k_{29}$ (Rate of Reaction 29) | 1 (per time) |
| Number of Wound Cells per new Cancer or TB cell ($n$) | 20 (wound cells / cancer or TB-infected cell) |
| Initial Value of $L_N$ | 0.5 (fraction hepatocytes) |
| Initial Value of $L_A$ | 0.5 (fraction hepatocytes) |
| Initial Value of $L_\phi$ | 0 (fraction hepatocytes) |
| Initial Value of $M_f$ | 100 (units muscle mass) |
| Initial Value of $M_e$ | 100 (units muscle mass) |
| Initial Value of $M_\phi$ | 0 (units muscle mass) |
| Initial Value of $M_{e\phi}$ | 0 (units muscle mass) |
| Initial Value of $W$ ($W_0$) | 1 if acute wound; $nC_0$ if cancer; $nTB_0$ if tuberculosis (units wound mass) |
| Initial Value of $C$ ($C_0$) | $10^{-4}$ (units wound mass) |
| Initial Value of $TB$ ($TB_0$) | $10^{-4}$ (units wound mass) |
| Initial Value of $A$ ($A_0$) | 1 (units concentration) |
| Initial Value of $\phi$ ($\phi_0$) | 0 (units concentration) |
| Initial Value of $P_I$ ($P_{I_0}$) | 0 (units concentration) |

### 2.4 Wasting Within the Model

There are two muscle wasting pathways that the body can use to obtain amino acids. The first, IMW, has been described above (Reactions 4 and 7). The second, homeostatic muscle proteolysis, relies on muscle as a reservoir of amino acids. If blood amino acid levels fall below the normal range, muscle can waste, releasing amino acids to supply the deficit (Reaction 4). Similarly, if blood amino acid levels are above the normal range, they can be incorporated into new muscle tissue (Reaction 5).

We incorporated both wasting pathways into our model to faithfully recapitulate physiology Therefore, when IMW is inhibited, the system attempts to use homeostatic muscle proteolysis to maintain amino acid levels in our simulations.

### 2.5 Dietary Intake Within the Model

Dietary intake is a state variable in our model of a biochemical reaction network. At each timestep, new nutrients flow into the system by using the following reaction equation:

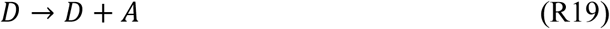

in which *D* is dietary intake, a state variable initialized at a value of 1. Patients with wounds often experience partial anorexia, the severity of which corresponds to the severity of the wound [2]. We model this phenomenon by allowing dietary intake and inflammation to interact, without adding another inflammation sink (Reaction 20).

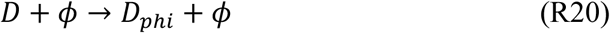

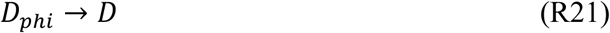

Dietary intake is reduced by inflammation until reaching a certain steady state value, which depends upon the amount of inflammation and the rates of Reactions 20 and 21. Then when the wound is resolved, dietary intake slowly returns to normal (Reaction 21). When incorporating therapeutic nutritional supplementation into our model, we set the reaction rates for Reaction 20 and Reaction 21 to 0. Consequently, *D* remains constant at the initial value.

Therefore, when we wanted to simulate nutritional support, we set the rates for Reactions 20 and 21 to 0, keeping the value of *D* constant, and initialized *D* at a value of 1. When we wanted to simulate nutritional support at 2× the normal dietary intake, we initialized *D* at a value of 2.

### 2.6 Metrics of Wound Healing (Figure 2)

### 2.7 Experiment to Evaluate Our Hypothesis

To evaluate if the absence of IMW impairs wound healing, we measured the time needed to heal an acute wound (such as a burn) over a continuous range of IMW.

**Figure 2.**
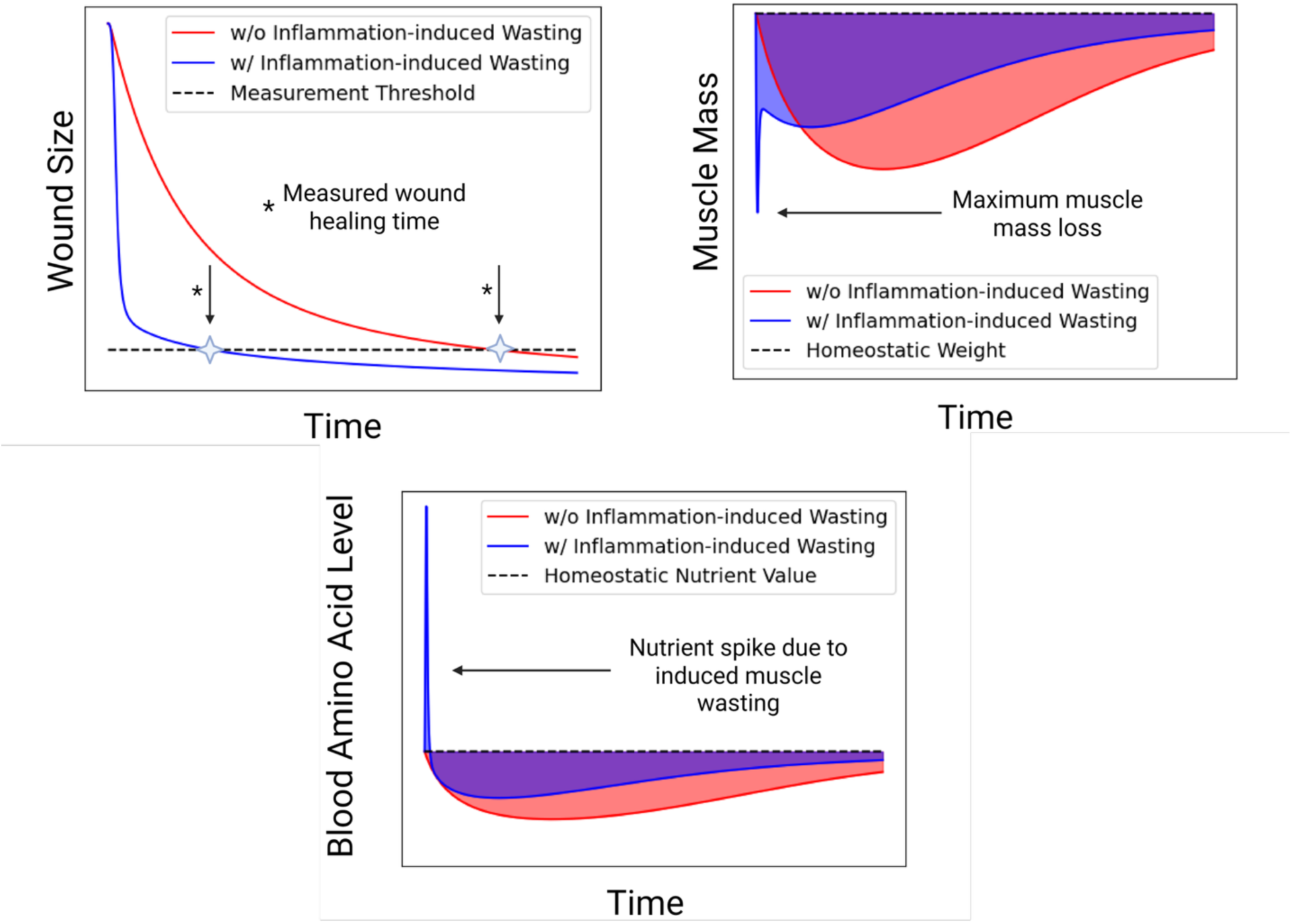
Metrics to measure wound-healing acceleration in the model’s simulations. The *wound-healing time* measures how long it takes the system to clear 90% of the wound. This enables quantification of the difference in wound healing caused by IMW. The *area under the blood amino-acid–level curve* shows the cumulative perturbation of amino acid levels caused by the wound. The *area under the muscle mass curve* gives a measure of the cumulative effect of the wound on muscle mass over the course of wound healing. The *maximum muscle mass loss* indicates the severity of wasting during wound healing.

To simulate the absence of IMW in the experiment, a new reaction equation was introduced in which uninflamed myocytes still interact with and provide a sink for inflammatory cues, but their state remains constant (Reactions 22-23).

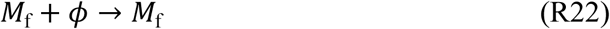

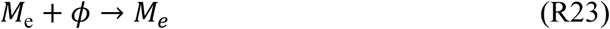

To test a continuous range of experimental scenarios between no IMW and full IMW, we simultaneously varied the reaction rate of these reactions and that of Reactions 7 and 8, always keeping their sum equal to 1. This is analogous to setting a probability, which we denote as *R*, that a myocyte will initiate the wasting program upon encountering an inflammatory cue. By simulating the system across *R* values ranging from 0 to 1, we treat IMW as a continuous independent variable in our acute wound healing experiments.

Knowing that IMW depends on the amount of inflammatory signals in the system, we also varied the rate of inflammation generation (Rate of Reaction 25: *k*_25_) to observe the resultant effects on wound healing and muscle loss. We tried three different values for *k*_25_: a “low” inflammation value – 4; a “medium” inflammation value – 8; and a “high” inflammation value – 12.

To simulate an acute wound, the wound compartment’s initial condition was set to 1. After simulating the system for 500 units of time, wound-healing metrics were calculated as shown in Figure 2, with results shown in Figure 3.

**Figure 3.**
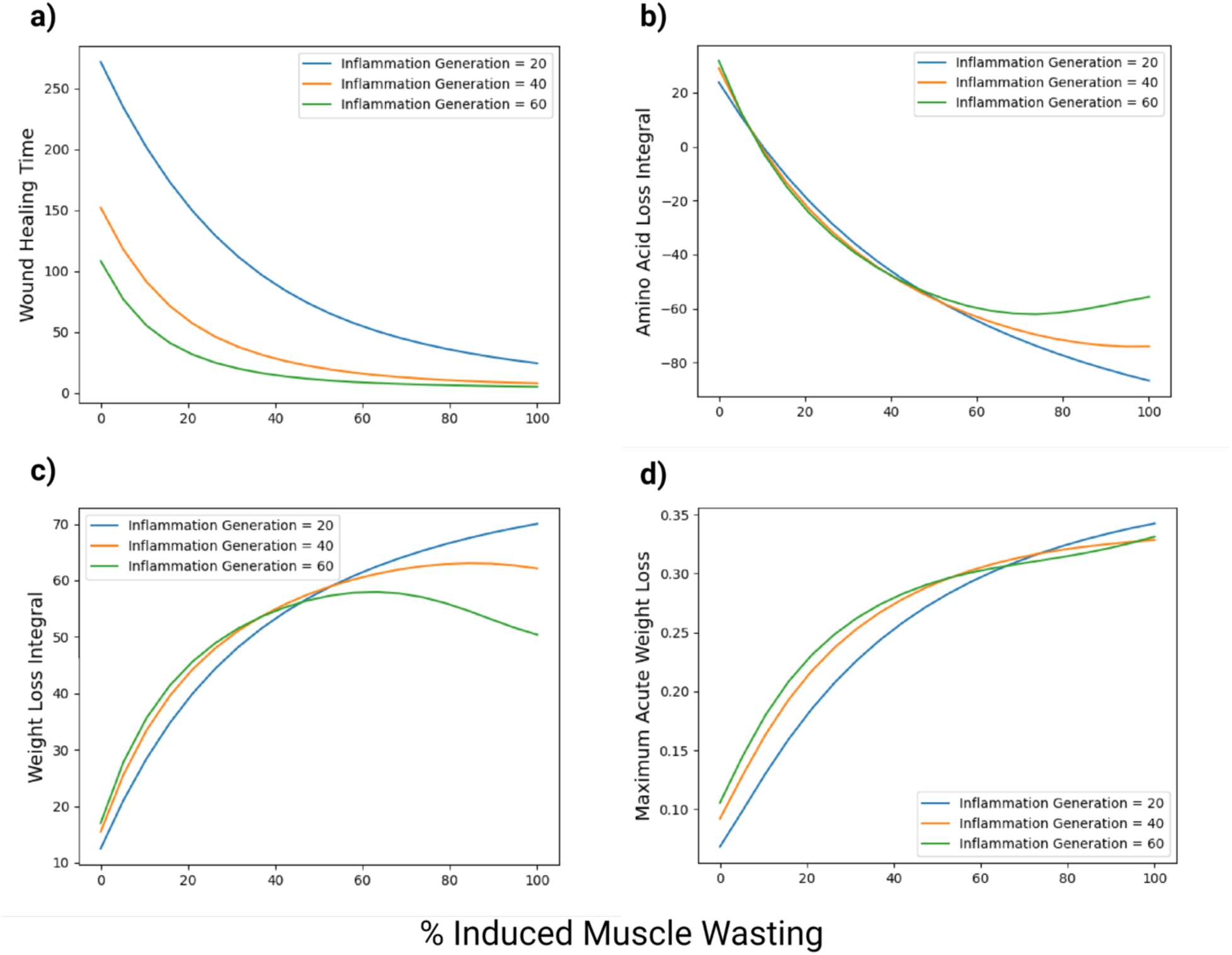
IMW improves the rate of wound healing. (a) Across all inflammation generation parameter values, the time needed for wound healing decreased with IMW. (b) Increased levels of IMW also decreased the cumulative perturbation to amino acid levels in the system. (c) However, as IMW increases, the time integral of muscle mass loss generally increases. (d) All the benefits of IMW come at the cost of increased acute muscle mass loss as lean mass is wasted.

**Figure 4.**
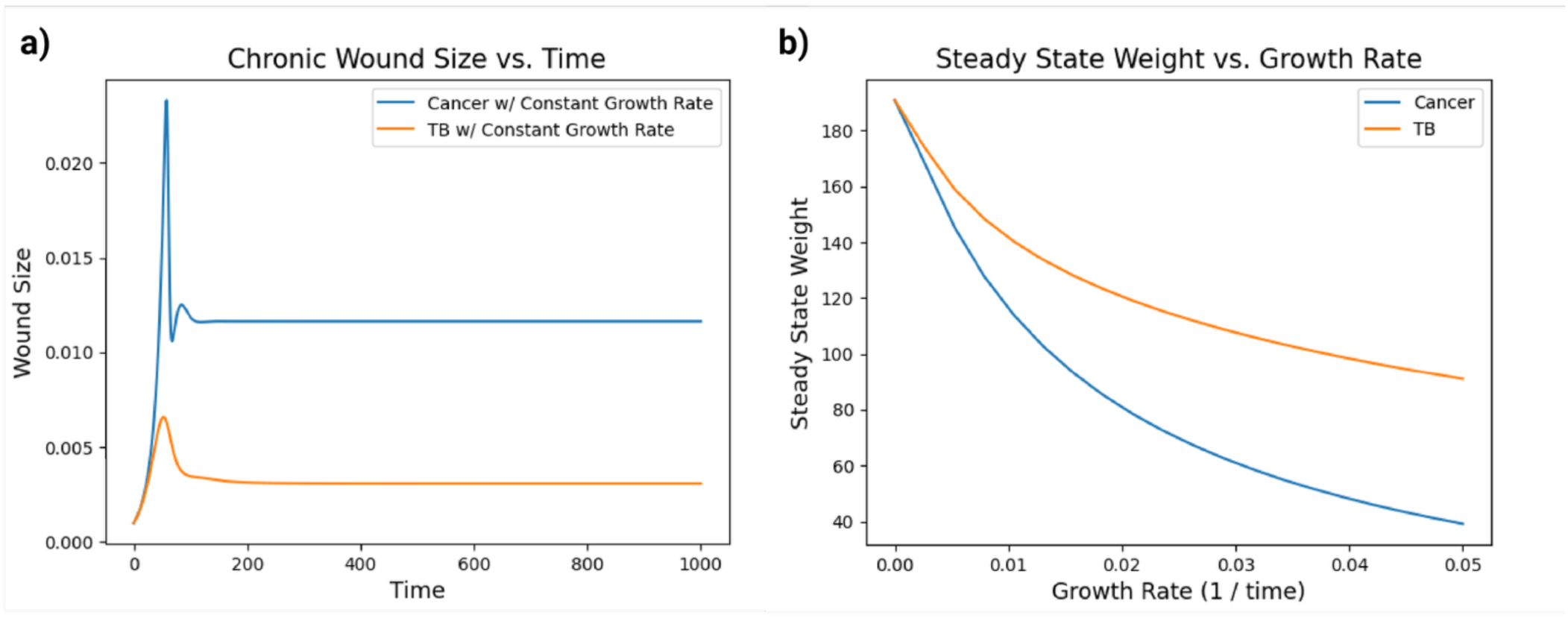
Increased pathogen growth rate causes increased wasting from chronic wounds. (a) At first, the cancer and tubercular mycobacterium populations experience exponential growth. As IMW begins, the spike in amino acid levels (and thereby APR activity) facilitates increased immune destruction of the populations, causing them to peak and return to a lower steady-state value, at which population growth is balanced by immune destruction. Note that the cancer population grows faster because of its ability to use the released amino acids to aid proliferation. (b) This bifurcation diagram shows how the steady state muscle mass depends on the growth rate of the pathogenic populations. The amount of wasting is minimal at low growth rates, but it rapidly becomes severe as the pathogens’ growth rate increases.

### 2.8 Effects of IMW on Chronic Wounds

To implement conditions such as cancer and TB in our model, we introduced reactions to allow the pathogenic agents to grow exponentially within the patient. For cancer, this reaction is given by

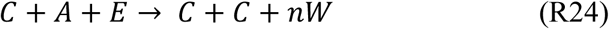

in which *W* is the wound compartment, and *n* is a parameter controlling how many noncancerous cells begin releasing inflammatory cues as the result of the increase in the cancer population.

This captures how a cancerous population mixes with noncancerous stroma as it grows, and in the process forms a tumor. Being human in origin, the cancer cell is assumed to use large quantities of bloodborne amino acids to replicate itself. Then, the cancer cells also give off inflammatory signals (Reaction 25) and can be destroyed upon encountering acute-phase proteins and the immune cells that interact with them (Reaction 26).

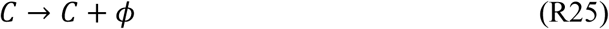

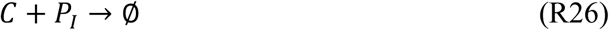

For TB, the growth reaction was

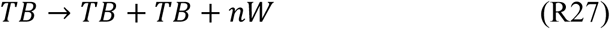

in which *TB* is an infected human host cell; *W* is the wound compartment; and *n* is a parameter controlling the number of nearby cells that begin giving off inflammatory cues as the result of increasing the infected host cell population. Because the number of infected host cells can grow exponentially, this process is not as straightforwardly dependent upon amino acids as in cancer. Therefore, we assumed that the growth kinetics for TB in our model would not be affected by amino acid levels. Because TB-infected cells emit inflammatory signals (Reaction 28) and are subject to immune destruction (Reaction 29), we also added reactions capturing these processes.

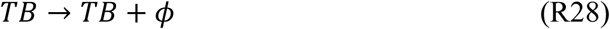

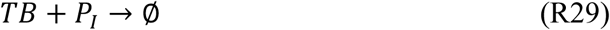

### 2.9 Effects of Nutritional Support on Wound Healing

To examine the effect of nutritional support on wound healing in our model, we ran simulations of both acute wounds and cancer with three possible dietary intakes: wound-induced anorexia– where the diet is actively suppressed by inflammatory signals–normal diet, and twice normal diet. Then, for acute wounds, we compared the wound-healing times and maximum acute mass-loss curves across these three diets.

Within the cancer context, we compared the relationship of the cancer-population size and the muscle mass reached at steady state with the growth rate. We varied both the dietary intake and level of IMW. These results were then plotted and are shown in Figure 5.

**Figure 5.**
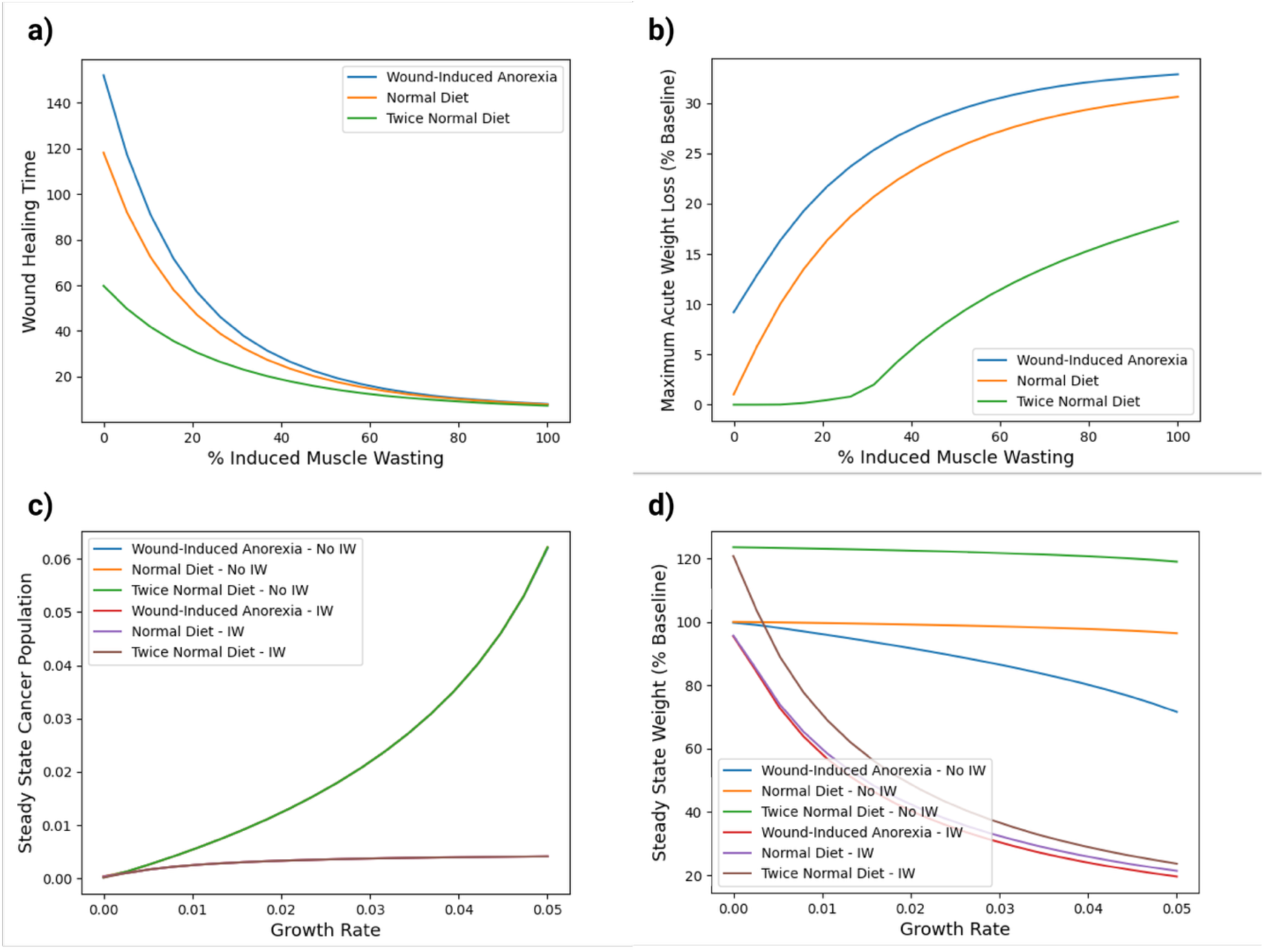
Nutritional support reverses cachexia when IMW is inhibited. (a) The influx of nutrients from nutritional support can accelerate wound healing, especially if nutrition intake is approximately twice what is normally consumed. Note that this is also true when IMW is inhibited. (b) Nutritional support can also decrease acute muscle mass loss associated with IMW. However, even when nutrient intake is twice the normal rate, acute muscle mass loss remains high (>10%) unless wasting is inhibited. (c) Dietary intake appears to have little consequence for the size of the cancer population at equilibrium, but the presence of IMW greatly inhibits the growth of cancer. (d) This graph summarizes why chronic wounds lead to cachexia that cannot be reversed with nutritional support. As the growth rate of the disease increases, the loss of mass from IMW increases as well, despite supranormal intake of nutrients. But when IMW is inhibited, then muscle mass is easily stabilized through nutritional support.

## 3. Results

Wound-healing time substantially decreases with IMW (Figure 3a). Not only do wounds heal faster in the presence of the muscle-wasting response, but the amino acid levels in the system were stabilized, reducing the strain of wound healing (Figure 3b). Additional amino acids released into the blood enhance the liver’s APR. However, the model correlates increased muscle mass loss over time with the level of wasting (Figure 3c). And while acute muscle mass loss also increases with IMW, simulations with high inflammation showed roughly the same acute muscle mass loss as those with low inflammation, demonstrating that enhanced induction of the muscle-wasting response can paradoxically preserve muscle mass by more rapid wound healing (Figure 3d).

After we observed the effects of IMW on healing an acute wound, we then considered how this affects a chronic wound, exemplified by cancer or a TB infection. In our model, these two pathogens are distinguished by how they can use resources within the body, as discussed above in Section 2.8. Both populations grew quickly before the IMW and APR were fully activated. Then, after an initial reduction in population, an equilibrium population size was reached in which proliferation equals immune destruction (Figure 4a).

We became curious about how the muscle mass of the patient would depend upon the proliferation rate of the pathogen, which is often correlated with virulence [17]. The steady-state muscle mass of the patient indeed decreased sharply with the increased growth rate of the pathogen, and when the growth rate was small, the patient’s muscle mass loss was also small (Figure 4b). Based on these results, our model would predict that a patient with progressive disease (that is, with an increasing pathogen growth rate) will experience progressive cachexia over time.

Finally, because nutritional support is such an integral part of cachexia care, we wanted to examine its effects on wound healing in our model, within both the acute-wound and cancer contexts. Because wound healing is directly related to the availability of nutrients, the speed of acute-wound healing was increased by providing stable dietary intake, especially in the absence of IMW (Figure 5a). But we observed that, despite an elevated intake of nutrients, acute muscle mass loss remained severe provided that IMW was still present (Figure 5b).

Within the cancer context, the production of immune proteins remained low without IMW. Therefore, as the growth rate of cancer was increased, the equilibrium size of the cancer population increased rapidly provided that IMW was absent. But in the presence of IMW, the cancer population reached only a fraction of the size it had with no IMW (Figure 5c). While IMW featured reduced cancer growth, it also caused dramatically more muscle mass loss as the cancer growth rate increased, for all dietary intakes. In contrast, without IMW, the patient’s muscle mass remains stable at baseline when nutritional supplementation is given (Figure 5d).

## 4. Discussion

We hypothesized that IMW is a mechanism that supports wound healing, and we tested this by performing an experiment within our model. The results of that analysis show that when IMW is inhibited, wound healing is drastically slower because of the slower synthesis of acute-phase proteins. In addition, the amino acid levels in the blood are prevented from falling dangerously low. In exchange for this boost to wound healing, the body suffers from acute muscle mass loss as muscle is rapidly catabolized (Figure 3).

Why is this tradeoff beneficial? First, we can observe that timely wound healing is essential for an organism to survive. If a wound remains open, it could become infected with micro-organisms, and the loss of heat, water, and nutrients to the environment might be fatal.

Furthermore, if an injured body part is not healed quickly, it may be injured again before it is repaired, leading to even greater impairment. Therefore, such a critical need is a worthy cause for expending some fat and muscle.

Instead of catabolizing bodily reserves, why shouldn’t the organism consume the needed nutrients from its environment? One immediate problem with this solution comes from the wound. If the damage to the body impairs seeking and obtaining food, as a severe fracture, burn, or infection does, then simply eating more is not feasible. And if a creature seeks to intake even more nutrients than normal, the amount of energy expended to acquire, digest, and process food will be counterproductive and detrimental. Using the nutrients already present in muscle is more efficient. In light of these observations, wound-induced anorexia makes sense as an adaptation to conserve energy and concentrate resources on resolving the wound. Indeed, the same signals that cause wound-induced anorexia can also induce wasting [18–19].

Other phenomena that occur alongside IMW can also be understood as an adaptive reshuffling of metabolic priorities. For example, patients displaying IMW often experience peripheral insulin resistance, due to modification of the insulin receptor in response to cytokines like TNF-α [20]. This modification sequesters glucose in the bloodstream, where it can fuel wound healing.

Because a profound fatigue accompanies these processes, patients often rest or use their muscles less than normal, reducing the muscles’ need for glucose during wound healing. The selective wasting of type IIa/x and type IIx glycolytic muscle fibers, another phenomenon broadly associated with cachexia, furthers this end [21]. Taken together, these mechanisms ensure a ready supply of nutrients for wound healing.

However, this ensemble of adaptations, which can save the life of a wounded patient when treatment is inaccessible, often hinders treating unresolved disease in hospital settings. While liberating more nutrients for wound healing, it may accelerate the growth of some disease-causing agents, such as cancer, although it is unclear if increased immune activity counteracts this effect (Figures 4a, 5c). Moreover, as unresolved disease grows worse and generates more inflammatory cues, IMW will grow worse, even with considerable dietary supplementation (Figure 5d). This explains why cachexia cannot be reversed by “conventional nutritional support” [1].

If nutritional support cannot reverse cachexia by itself, then how can it be treated? The model predicts that once IMW is inhibited, nutritional support can stabilize a patient’s muscle mass over all disease growth rates (Figure 5d). Therefore, if a reliable method for inhibiting IMW is found, cachexia can be cured.

The current literature on inhibiting IMW describes two different approaches: dietary and pharmacological. There is evidence that muscle and fat may partially cease wasting in response to dietary increases in certain nutrients, and patients with TB often regain lost fat mass, if not muscle mass, during antimycobacterial therapy with nutritional support [22–23]. However, it is unclear if this is caused by increased endocrine, anabolic signaling, or decreases in breakdown at the cellular level. Further research is needed to clarify how wasting pathways can be regulated by nutrients.

Pharmacological approaches, in contrast, aim to directly block wasting pathways within muscle and fat cells. For example, inhibiting myocytes’ expression of MuRF1—an E3 ubiquitin ligase responsible for degrading many proteins related to muscle contraction—may be possible [24]. These methods seem promising; however, our model indicates that they may not succeed in treating cachexia without joint administration of nutritional support. While causing muscle mass loss, IMW provides essential nutrients for wound healing. If IMW is inhibited, nutritional support will prevent dangerous drops in amino acid blood levels. In this manner, the combination of IMW inhibition and nutritional intervention may resolve patients’ cachexia, enabling physicians to concentrate on healing other aspects of their disease.

There are limitations to our approach. Our mathematical model, while including the most salient participants in IMW, does not capture the full complexity of the human body. There are other systems that play an integral role in cachexia’s development: adipose tissue, the central nervous system, etc. Notwithstanding the increased complexity of a real body, our model provides a starting point for analysis of cachexia by showing what results from these simple interactions without confounding factors. Another limitation of our study is that our model was not fit to data from patients or *in vivo* models. This is chiefly because few studies, if any, have measured the metrics that our model outputs, and therefore, there are no comprehensive datasets on which to fit our model. But our model qualitatively matches the typical course of muscle mass loss seen after severe burns, giving reasons to be confident in its accuracy [16]. Additionally, we plan to complete future studies in which we will measure all the data needed to evaluate the model and its conclusions in an *in vivo* setting.

Cachexia presents great challenges to patient care, reducing tolerance to therapies, quality of life, and survival. Decades after the first studies of its molecular mechanisms, elucidating cachexia’s etiology is more important than ever [7]. The success of future treatment depends on it. In pursuit of this objective, we presented a novel paradigm of cachexia, hypothesizing that induced muscle wasting is an adaptation to support the material needs of rapid wound healing. To evaluate this, we developed a mathematical model of cachexia, deriving immediate implications for treating patients. To mitigate and reverse cachexia, nutritional support alone is not sufficient. Nutritional interventions will only succeed when induced muscle wasting has been properly inhibited.

However, *once wasting has been turned off*, maintaining stable muscle mass becomes comparatively easy. We eagerly anticipate the discovery of improved methods to inhibit induced muscle wasting, relieving sufferers of the consuming grip of cachexia.

## Acknowledgments

Editorial assistance was provided by Gerard Hebert of Moffitt Cancer Center’s Biomedical Research Library; no compensation was given beyond his regular salary. This work was funded by an NIH U54 grant (U54CA274507). Biorender was used in the creation of figures.

## Ethical Standards

The authors assert and certify that they have complied with the Ethical Guidelines specified by the *Journal of Cachexia, Sarcopenia, and Muscle Wasting*. The manuscript does not contain clinical studies or patient data. No animal or human research was conducted as part of this study.

## Data and Code Availability

The code used to run simulations and generate plots for this manuscript can be found in the following GitHub repository: https://github.com/chaynes2019/Wasting-is-essential

